# RNANetMotif: identifying sequence-structure RNA network motifs in RNA-protein binding sites

**DOI:** 10.1101/2021.09.15.460452

**Authors:** Hongli Ma, Han Wen, Zhiyuan Xue, Guojun Li, Zhaolei Zhang

## Abstract

RNA molecules can adopt stable secondary and tertiary structures, which is essential in mediating physical interactions with other partners such as RNA binding proteins (RBPs) and in carrying out their cellular functions. In vivo and in vitro experiments such as RNAcompete and eCLIP have revealed in vitro binding preferences of RBPs to RNA oligomers and in vivo binding sites in cells. Analysis of these binding data showed that the structure properties of the RNAs in these binding sites are important determinants of the binding events; however, it has been a challenge to incorporate the structure information into an interpretable model. Here we describe a new approach, RNANetMotif, which takes predicted secondary structure of thousands of RNA sequences bound by an RBP as input and uses a graph theory approach to recognize enriched subgraphs. These enriched subgraphs are in essence shared sequence-structure elements that are important in RBP-RNA binding. To validate our approach, we performed RNA structure modeling via discrete molecular dynamics folding simulations for selected 4 RBPs, and RNA-protein docking for LIN28. The simulation results, e.g., solvent accessibility and energetics, further support the biological relevance of the discovered network subgraphs.

**Author Summary:** RNA binding proteins (RBPs) regulate every aspect of RNA biology, including splicing, translation, transportation, and degradation. High-throughput technologies such as eCLIP have identified thousands of binding sites for a given RBP throughout the genome. It has been shown by earlier studies that, in addition to nucleotide sequences, the structure and conformation of RNAs also play important role in RBP-RNA interactions. Analogous to protein-protein interactions or protein-DNA interactions, it is likely that there exist intrinsic sequence-structure motifs common to these RNAs that underlie their binding specificity to specific RBPs. It is known that RNAs form energetically favorable secondary structures, which can be represented as a graph, with nucleotides being nodes and backbone covalent bonds and base-pairing hydrogen bonds representing edges. We hypothesize that these graphs can be mined by graph theory approaches to identify sequence-structure motifs as enriched sub-graphs. In this article, we described the details of this approach, termed RNANetMotif and associated new concepts, namely EKS (Extended K-mer Subgraphs) and GraphK graph search algorithm. To test the utility of our approach, we conducted 3D structure modeling of selected RNA sequences through molecular dynamics (MD) folding simulation and evaluated the significance of the discovered RNA motifs by comparing their spatial exposure with other regions on the RNA. We believe that this approach has the novelty of treating the RNA sequence as a graph and RBP binding sites as enriched subgraph, which has broader applications beyond RBP-RNA interactions.

## 1. INTRODUCTION

The human genome encodes approximately 1500 or more RNA binding proteins (RBPs), which regulates every aspect of the RNA biogenesis and RNA biology, including RNA splicing, modification, degradation, protein translation, and RNA subcellular localization (1-6). RBPs are also involved in many important developmental processes such as embryogenesis, proliferation, and differentiation. Mutations or dysregulation of RBPs are also implicated in many human diseases including cancer (7-9). In vitro methods such as RNAcompete, RNA Bind-n-Seq (RBNS) and high-throughput RNA-SELEX (HTR-SELEX) can measure binding affinity of an RBP to RNA oligomers (10-12), and in vivo methods such as PAR-CLIP, iCLIP, and eCLIP can identify regions on the RNA transcript that are bound by RBPs in cells (13-15). Analysis of these in vitro binding affinity data and in vivo binding sites have revealed that the structure of RNA molecules can help understand the binding mode between a specific RBP and their target sequences. For example, RNA binding domains can be grouped into single stranded or double stranded RNA binding domains, ssRBD and dsRBD, based on their preference for RNA targets that are either single stranded (unpaired) or double stranded (paired). A number of computational methods had been developed to ascertain the structure properties of these RBP bound RNAs with the aim to incorporate structural information into a predictive model that can help decode mechanisms that regulate protein-RNA interactions (16-33). A number of these representative methods are briefly described below.

The structure-based methods differ in how they encode RNA structure information in their models thus they can be grouped into two broad categories according to the abstract level of structural encoding. Methods like MEMERIS (18), mCarts (29) and SMARTIV (22) apply simplified structure representations, including RNA accessibility score and discrete secondary structure notations such as paired and unpaired, while methods including RNAcontext (17), ssHMM (24), BEAM (25), GraphProt (16) and SARNAclust (27) characterize the shape of substructures with different labeling methods. Approaches such as RNAcontext (17) explicitly label individual nucleotides with an additional feature indicating the secondary structure of the particular nucleotide, i.e., paired, hairpin loop. Methods such as Graphprot (16) and ssHMM (24) further captured structure inter-dependencies between neighboring nucleotides through graph based encoding or hidden Markov model. Moreover, methods such as BEAM (25) attempted to explicitly encode the pattern of nucleotide base pairing and catalogue the occurrence of specific secondary structure motifs based on the length of the stem or the size of the hairpin loop. SARNAclust (27) comprehensively studied the topology and annotations of the complete RNA structure and provides several options for graph transformations of RNA shapes. Despite these advances, our understanding on the structural basis of binding events between protein and RNA still lags our understanding on the interaction between protein partners, which is largely due to the lack of atomic resolution RNA structures.

In parallel to the study of protein-RNA interactions, our understanding on protein-DNA interactions has been greatly aided by the availability of a plethora of structure information on protein-DNA complexes and on DNA structures alone. It has been recognized that the overall shapes of the DNA molecules play an important role in the initial recognition by proteins and the subsequent binding processes (34,35). These structural elements allow optimal positioning between amino acids and the interacting moieties on DNA molecules, e.g., backbone or nucleosides. It is likely that similar principles are also important in protein-RNA recognition, i.e., there exist recurring RNA 2-D or 3-D structure motifs that are essential in the stabilization of the RNA molecule and presenting the nucleotide moieties to RBPs (36-39). The 2-D structure of an RNA molecule consists of a network of base-pairing interactions between nucleotides, which, without atomic resolution structure, is the best approximation of a RNA’s 3-D structure. We hypothesize that there exist intrinsic subgraphs in these networks that can potentially separate RNA sequences that are bound by a specific RBP or RBPs from those that are not bound; we term these 2-D sequence entities as RNA network motifs.

Interactions between RNA-binding proteins (RBPs) and RNA molecules employ diverse and dynamic modes (40), finding the exact structural motifs has been a challenging problem. There have been previous attempts trying to solve this problem in an approximate way (16,19,24,25). Despite the recent advances, there remain two issues that need to be addressed. First, we need to combine sequence and structural information of RNA in an efficient and flexible way. Global approaches such as GraphProt (16) model RNA structure as complete graphs; other methods such as BEAM (25) use an additional alphabet to represent the structural state of each nucleotide. A framework that can both capture the states of individual nucleotides while encoding the base-pairing information between the nucleotides would be more desired. Second, given that RNA structures are represented as 2-D or 3-D graphs, we need efficient and robust approaches to search many of these input structures and identify enriched and potentially discriminative network motifs.

Towards these goals, we herein introduce a novel algorithm, RNANetMotif, which takes as input thousands of RNA sequences presumably bound by an RBP and uses a graph theory approach to search for “base pair derived subnetworks” enriched in the predicted RNA secondary structures. RNANetMotif consists of the following steps. (i) For each RNA locus bound by an RBP as determined via eCLIP experiments, we predict its base-pairing pattern using software RNAplfold and represent its secondary structure as a network. (41). (ii) We developed a novel GraphK algorithm, which can break apart the aforementioned RNA secondary structure network into subgraphs consisted of both RNA backbone and base pair interactions. For each RBP, these subgraphs are pooled and filtered to obtain a candidate pool. (iii) We compute HVDM (Heterogeneous Value Difference Metric) distance matrix to construct a similarity network with candidate subgraphs being nodes, and further conducted network pruning. (iv) In the similarity network, we next perform maximal clique enumeration and rank the occurrence of nodes in large maximal cliques to obtain enriched subgraph candidates, i.e., RNA net motifs. (v) Finally, we conducted 3D structure modeling of selected RNA sequences through discrete molecular dynamics (MD) folding simulation and evaluated the significance of the discovered network motifs by comparing their spatial exposure with other regions (42).

We note, in addition to RBP binding sites, RNANetMotif can also be applied in other scenarios to extract enriched RNA sequence or structure motifs. RNANetMotif compares favorably with other methods in terms of performance and run-time. We believe RNANetMotif offers a fresh approach in understanding the interactions between RBP and RNA targets, and in extracting informative RNA motifs. To make this tool more accessible to the community, we constructed a web server (http://rnanetmotif.ccbr.utoronto.ca), that stores the results from our analysis of ENCODE RBP datasets (16 RBPs) and allows users to upload a collection of RNA sequences of their own interests for motif analysis. The RNANetMotif is a new way of investigating RNA sequences and can be further extended into analysis of other categories of RNA sequences.

## 2. MATERIALS AND METHODS

### 2.1 Collection of RBP eCLIP data and preprocessing

We downloaded the “enhanced UV crosslinking followed by immunoprecipitation” (eCLIP) data (13) from ENCODE project website (release v101) (https://www.encodeproject.org/) in June 2020, which consists of chromosomal peak regions that are bound by specific RBPs. The peaks were annotated as IDR BED files which had been processed following the eCLIP-seq Processing Pipeline. Reproducible and significant peaks that passed the Irreproducible Discovery Rate (IDR) were identified by a modified IDR method (1).

A total of 223 IDR files corresponding to 150 unique RNA binding proteins (RBPs) were collected, which comprised of 120 files for K562 cell line, 103 files for HepG2 cell line. We next retained only 56 IDR files that have more than 5,000 mapped binding sites. We further excluded those proteins that were annotated to be primarily involved in mRNA splicing and excluded those RBPs that had fewer than half of the binding sites mapped to the annotated exon regions. The final derived set contained 22 IDR files. The rational of above procedures is to limit our study only to those RBPs that primarily bind to the mature mRNA transcript region. To unambiguously extract exon regions, we used the most prominent transcript for each gene as was defined based on the basic gene annotation from GENCODE (Release 34, GRCh38.p13) through hierarchical filtering: first filtered by APPRIS annotation (highest priority) (43), then by transcript support level, and finally by transcript length (longer isoform preferred). We only kept one prominent transcript per gene.

We next processed the 22 IDR files by only keeping the peaks that are entirely localized in one of the exons of the prominent transcript., i.e., removing those binding sites in the introns. The detailed information of the 22 RBPs with specific cell line is summarized in **Table 1**. We chose 100 nt as the uniform binding site length (50 nt extensions up- and down-stream of the center position) and used ‘getfasta’ function from ‘bedtools’ to map BED files to FASTA files. We then removed redundant and missing data in the binding sites by using module ‘cd-hit-est’ from CD-HIT (44,45) at 80% similarity cut-off.

**Table 1.** RBPs and the dataset information.

| RBP and Cell Line | RNA binding domain | #peaks |
| --- | --- | --- |
| AKAP1 HepG2 | KH,Tudor | 5338 |
| BCLAF1 HepG2 |  | 22884 |
| DDX24 K562 | Helicase ATP-binding,Helicase C-terminal | 5841 |
| DDX3X HepG2 | Helicase ATP-binding,Helicase C-terminal | 5976 |
| DDX3X K562 | Helicase ATP-binding,Helicase C-terminal | 3961 |
| FAM120A K562 |  | 4218 |
| G3BP1 HepG2 | NTF2, RRM | 5204 |
| GRWD1 HepG2 | WD1, WD2, WD3, WD4, WD5, WD6, WD7 | 16040 |
| IGF2BP1 K562 | RRM1, RRM2, KH1, KH2, KH3, KH4 | 4690 |
| LARP4 HepG2 | HTH La-type RNA-binding,RRM | 4350 |
| LIN28B K562 | CSD | 4616 |
| PABPC4 K562 | RRM1, RRM2, RRM3, RRM4, PABC | 4865 |
| PPIG HepG2 | PPIase cyclophilin-type | 13538 |
| PUM2 K562 | PUM-HD, Pumilio 1, Pumilio 2, Pumilio 3, Pumilio 4, Pumilio 5, Pumilio 6, Pumilio 7, Pumilio 8 | 4742 |
| RBM15 K562 | RRM1, RRM2, RRM3, SPOC | 6800 |
| RPS3 HepG2 | KH type-2 | 4697 |
| SND1 HepG2 | TNase-like 1, TNase-like 2, TNase-like 3, TNase-like 4, Tudor | 5697 |
| UCHL5 K562 |  | 12866 |
| UPF1 HepG2 |  | 8547 |
| UPF1 K562 |  | 11708 |
| YBX3 K562 | CSD | 12706 |
| ZNF622 K562 |  | 10551 |

### 2.2 Overall workflow of the RNANetMotif method

As shown in **Figure 1**, RNANetMotif consists of the following steps. In Step 1, we predict RNA base-pairing probability using RNAplfold and represent each RNA binding site as a graph in which edges represent base pairings and backbones. In Step 2, we developed a GraphK algorithm to partition the network into Extended K-mer Subgraphs (EKSes) and filter to obtain the final EKS pool. In Step 3, we compute the HVDM (Heterogeneous Value Difference Metric) distance matrix (46) and use network pruning to construct a similarity network among the EKSes. In Step 4, we identify overlapping densely connected modules on the similarity network and evaluate the significance of these modules by comparing them with a previously established negative set. We next identify the representative EKSes by conducting maximal clique enumeration, followed by extracting the overlapping large size cliques in the similarity network. In Step 5, for several RBPs, we conducted 3D structure modeling on the RNAs to further examine the network motifs in a structure and dynamic context. As an example, we selected LIN28B protein and performed protein-RNA docking to examine the detailed binding mode and evaluated the robustness of the RNANetMotif method.

**Figure 1.**
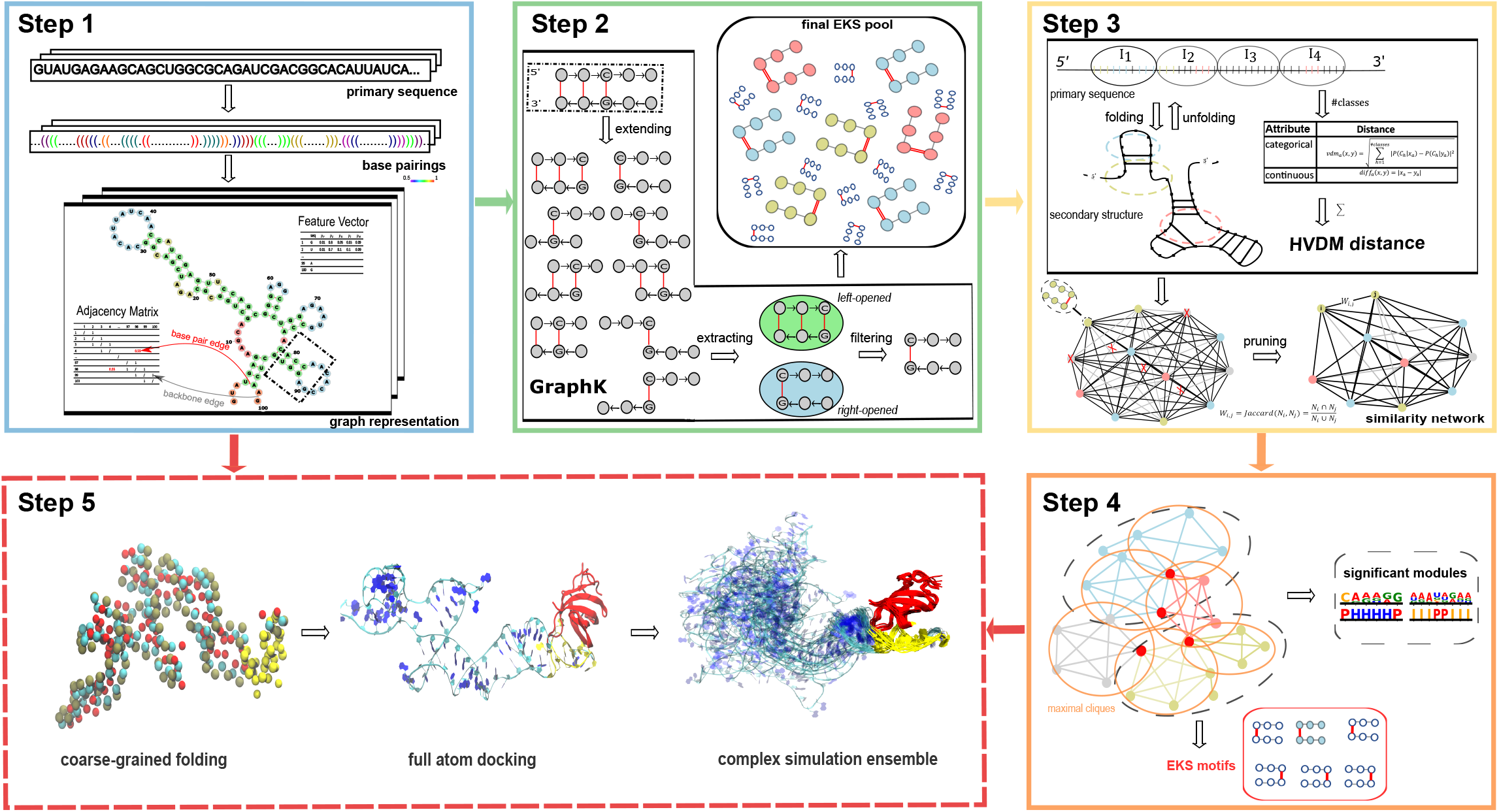
Workflow of the RNANetMotif method. **Step1**. Predicting base-pairings of protein-bound RNA sequences and graph representation. The representation includes nucleotide information (vertex feature vector) and base-pairing information (adjacency matrix), here depicted with different colors to distinguish backbone links (grey) from base pairing (red). **Step 2**. GraphK partition algorithm (see Methods) to obtain final EKS pool. **Step 3.** Calculate HVDM (Heterogeneous Value Difference Metric) distance matrix and construct similarity network of EKSes. We display the folding and unfolding between sequence and secondary structure to describe class choices to deal with categorical attributes in HVDM distance. After unfolding, EKSes (marked by dashed circles in the secondary structure) backed to primary sequence as gapped sequences with two k-mers falling into the four intervals {*Ik,k* =1,2,3,4} corresponding to the coordinates. **Step 4**. Detect significant network modules and then identify intrinsic EKS motifs. **Step 5**. RNA 3D structure modeling via discrete molecular dynamics folding simulations and protein-RNA docking with simulation for validation.

### 2.3 Predicting RNA secondary structures and graph construction

We used the RNAplfold program in the ViennaRNA package (version 2.4.13) for secondary structure prediction. The stem candidates were derived from the base pairing probability matrix calculated by McCaskill’s algorithm (41). After experimenting with the choices of window sizes (L and W-L), we set the parameters of RNAplfold as W=100, L=100 while allowing Watson-Crick (A:U and G:C) and wobble G:U base pairing (47). We note that, instead of generating a final structure, RNAplfold calculates base-pairing probabilities between nucleotide pairs in the entire sequence. For each RNA sequence, we only keep those reliable base pairs with probability >0.5, which do not form a pseudoknot and are in a centroid (48).

Having predicted base-pairing probabilities in the entire RNA, we next predicted and refined local secondary structure probabilities associated with each nucleotide by using the software RCK (20). The following five type representations are considered: H for hairpin loop, I for internal loop, M for multi-loop, E for external loop, and P for paired. The following parameters (W=100, L=100, u=1) were used. The final combined features for each nucleotide consist of its nucleotide type and the probabilities of having each of the five secondary structures, i.e., H, I, M, E and P.

We represented each 100 nucleotides (nt) long RBP bound RNA sequence as a weighted graph G = (V, E, w), where V consists of nucleotides encoded as vertices with discrete labels (A, C, G, U) and continuous labels (see above). The edge set E contains RNA backbones and the predicted base-pairings. The weight of backbone edges is defined as 1, while base-pair edges (Watson-Crick or G: U) as the probability predicted from RNAplfold.

### 2.4 Definition of EKS – Extended K-mer Subgraph

In this work, we introduce a new concept dealing with graph representation of local RNA secondary structures, referred to as Extended K-mer Subgraph or EKS. Given two base-pairing nucleotides, *i* and *j*, we build a subgraph by extending along the backbone edges from *i* and from *j*, separately and respectively (**Figure 2**). During the extension process, we traverse along the backbone from *i* and from *j* respectively, until a k-mer is reached in both directions. We define the resultant subgraph containing the *2k* number of nucleotides as an <u>E</u>xtended <u>k</u>-mer <u>S</u>ubgraph (EKS). The intuition of EKS is that these subgraphs represent smaller tractable network elements as component of a larger complex network.

**Figure 2.**
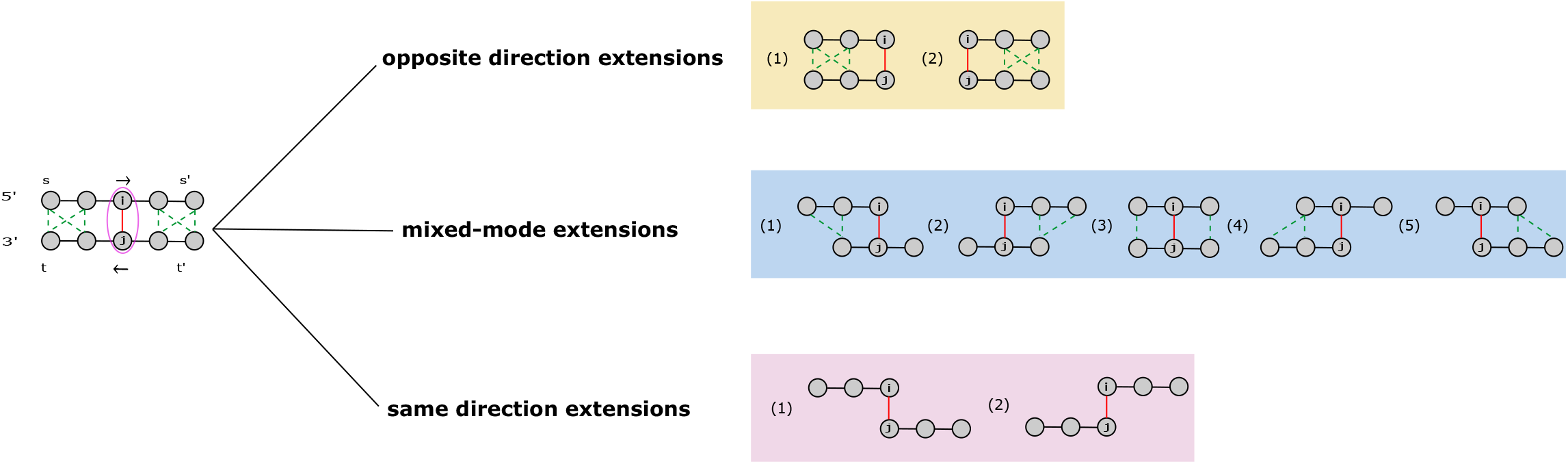
Definition and classification of EKS. As displayed, there are three categories of EKS: opposite direction extensions, mixed-mode extensions and same direction extensions. Black edge is the backbone edges, the red edge corresponds to the base pair interaction between i and j, and green dashed edge is the possible base pair interactions (there may be one or two of them appear, in most cases, only one of them exists in every graph for pseudoknot free RNA structure).

As illustrated in **Figure 2**, the linear sequence of the local folded RNA follows the following direction: 5′ → s → s′ → t′ → t → 3′. As an example, given k=3, we select base pair (i, j) as a root base pair, and denote U = {i, j} as the initial vertex set. We next perform k-mer extension process along the backbone edges, and extend U by adding nodes from the k-mer containing node i and the k-mer containing node j. After this process, the number of nodes in U reaches 2k and we denote it as G[U], the subgraph of G induced by U, as an EKS. The notation G[U] represents the subgraph of G with vertex set U and all the weighted edges that have both end vertices in U. We define the diameter of the graph as the “longest shortest path” between any two vertices in the graph. We also define the “compactness score “of G[U] as the reciprocal of the diameter of G[U], which is calculated by Floyd-Warshall algorithm with the time complexity of *O*(|*U*|^3^) (49,50). The benefit of using compactness score is that it falls in the range of (0, 1) thus is easier to manipulate in scoring or probability functions. Lower compactness score is indicative of lower “connectiveness” of the network, i.e., existence of node pairs separated by longer path.

Depending on the directions of traversing from the root base pair (i, j), we can divide the set of EKSes into three categories: opposite direction extensions, mixed-mode extensions, and same direction extensions. As shown at the top of **Figure 2**, the opposite direction extensions can generate EKSes that are either left-opened or right-opened subgraphs. Mixed-mode extensions have the most diverse shapes and topology. As for the same direction extensions, the network is extended in the same direction, both toward 3’ end or 5’ end. The nucleotides in these EKSes have lower chance of forming base-pairing unless pseudoknot structure is allowed.

### 2.5 GraphK – a novel subgraph extraction approach

Having defined EKS, we next introduce an algorithm, GraphK, which applies extension, extraction, and filtering steps to break a complex graph into overlapping representative EKSes. The detailed steps of the GraphK algorithm are as follows (**Figure 1 (Step 2)**). In Step 1, we conduct k-mer extension as described above in each RBP-bound RNA sequence to obtain all EKSes associated with the RNA sequence. In addition, we added an annotation for each EKS, including its type, compactness score, and the starting positions of the two k-mers. In **Step 2**, we extract EKSes generated from opposite direction extensions (i.e., right-opened or left-opened), and exclude those generated by opposite direction and mixed mode extensions. The rationale is to extract only those likely to form single stranded RNAs (see above). In **Step 3**, we filter EKSes by their compactness scores and only retain those with lower compactness scores, i.e., those with less base pairings. We next filtered the EKSes by the position of the nucleotides and kept the EKSes whose sequence is entirely located in the central 40 nucleotides. By this approach, we partitioned every 100 nt RNA into local network elements, further filtered and pooled them into the final EKS pool.

In the following, we elaborate the rationales behind the GraphK algorithm. First, it is difficult to represent the 2-D or 3-D structure of the entire RBP binding region. Therefore, we treated the predicted secondary structure of the entire RBP binding region as a network and extracted base-pair derived subgraphs from the network to represent components of the entire structure network. These local subgraphs are more tractable than a global graph in representing the entire RNA region. Our hypothesis is that the RBP bound RNAs could share some of these common subgraphs. Secondly, we briefly explain the choice of size, shape, and compactness score of the EKS. Since most single RNA binding domains (RBDs) appear to bind RNA motifs 3 ∼ 8 nt long (11), we chose *k* ranging from 3 to 5, corresponding to 2k ranging from 6 to 10. As a majority of the RBPs we studied prefer single stranded RNA, we only considered the subgraphs created by opposite direction expansion which generates single stranded RNAs (see **Figure 2**). In addition, we used a graph theory method to calculate compactness scores for EKSes and retained those with lower compactness score, i.e. those EKSes with sparse base-pairs and are more prone to binding by RBPs. Thirdly, we explain the choice of 40 nucleotides as the length of EKS. This is because the mean length of the peaks from eCLIP datasets is around 40 ∼ 60 nt and the structural distribution in the central regions of peaks shows better structural conservation.

### 2.6 Calculate HVDM distance matrix and construct EKS similarity network

We next calculate similarities between any pairs of EKSes so that we can select representative and discriminative EKSes associated with each RBP binding region. Each instance of EKS consists of 2*k* nucleotides and has 12*k* mixed-type attributes, comprised of 2*k* categorical attributes (i.e., nucleotide types as A, C, G, U) and 10*k* continuous attributes (i.e., secondary structure probabilities). We chose to use the heterogeneous value difference metric (HVDM) as the distance function, which uses value difference metric (VDM) to handle categorical attributes, and absolute differences to handle continuous attributes (46,51). Given two EKSes with the feature vector 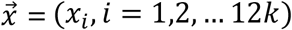 and 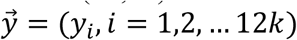, in the following we describe the details on the calculation of HVDM distance.

First, we classify an EKS according to which region on the RNA sequence the first and last nucleotide of the EKS falls into and assign the EKS to one of the following classes: *classes* = {*C*_*i*_, *i* = 1,2,3, …, 10}. As shown in **Figure 1 (Step 3)**, we first partition the central 40 nucleotides into 4 intervals of equal length of 10 nt as {*I*_*k*_, *k* = 1,2,3,4} and assign the EKS to one of the following classes according to the intervals into which the first and the last of the nucleotide falls: *C*_*i*_ ∈ {*I*_*i,j*_, *i, j* = 1,2,3,4. *i* ≤ *j*}. The VDM distance between 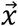 and 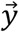 is defined in Equation (1), which considers the correlations between each possible attribute value and each EKS class.

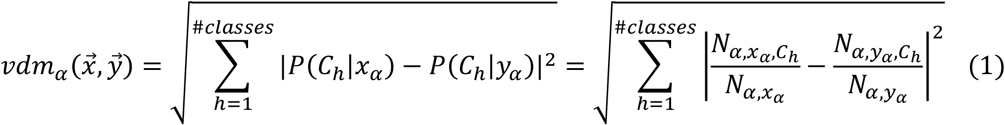

where

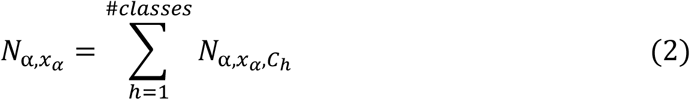

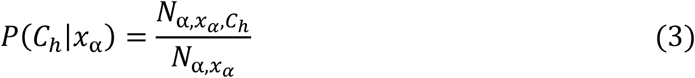

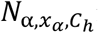 is the number of instances in all EKSes that have value *x*_*α*_ for attribute α and class *C*_*h*_; *N*_*α,x*_ is the number of instances in all EKSes that have value *x*_*α*_ for attribute α, i.e., 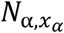 is the sum of 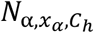 over all classes. *P*(*C*_*h*_|*x*_*α*_) is the conditional probability that the class is *C*_*h*_ given that attribute α has the value *x*_*α*_. The sum of *P*(*C*_*h*_|*x*_*α*_) over all classes is 1 for a fixed value of α and *x*_*α*_.

We next define the difference between two values *x*_β_ and *y*_β_ of a continuous attribute β as the absolute difference:

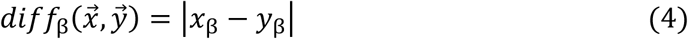

Taking these together, we then define the HVDM distance between 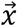 and 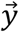 as:

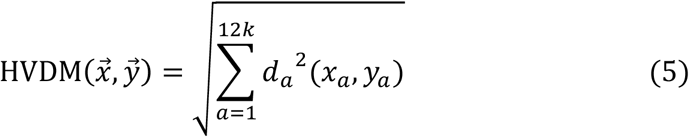

where

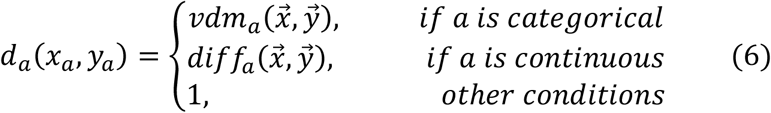

Next, we initialized a weighted complete graph S (*V*_*S*_, *E*_*S*_, *W*_*S*_) with EKSes as vertices and the above calculated HVDM distance between vertices as the weight of the edges. To denoise this graph, we then refined this graph by the following pruning steps. Firstly, we selected those vertices that are close to each other and have similar partners on the network. For *v*_1_, *v*_2_ ∈ *V*_*S*_, if *v*_1_ and *v*_2_ are among top n nearest neighbors of each other, i.e., *v*_1_ is one of the top n nearest neighbors of *v*_2_ and *v*_2_ is also one of the top n nearest neighbors of *v*_1_, *v*_1_, *v*_2_ and the edge between them (i.e., 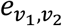) are retained. We then removed the remaining vertices and edges. By doing so, we obtained a vertex-pruned network and each pair of nodes in it is in a very close relationship with a short HVDM distance. To reveal the relationship between vertices in the vertex-pruned network more discriminatively, we replaced the distance-based similarity with neighborhood-based similarity and re-defined the weight of the edges 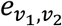 between nodes *v*_1_ and *v*_2_ in this network as the Jaccard index in Equation (7). 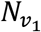 and 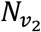 are the neighbor sets of *v*_1_ and *v*_2_ respectively.

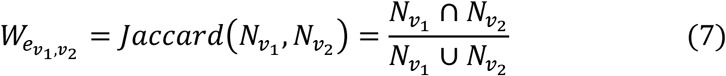

In the second pruning step, we removed any edges e with weight lower than a cutoff, *w*_*e*_ ≤ *w*_*cutoff*_; the purpose of this was to retain only those sufficiently similar EKS candidates. After experimenting with different parameters, we set n= 2% × *size of V*_*S*_, w_cutoff_ = 0.9 *quantile of all weights*. We denote the pruned graph as the similarity network among filtered EKSes. In **Supplementary Table 1**, we summarize the number of the nodes and edges of the similarity network for the 22 RBPs.

### 2.7 Overall binding preference and intrinsic network binding motifs

Having constructed the similarity network among EKSes for each RBP, we next took a multi-steps approach to identify EKSes that are important in RBP binding. We first searched the similarity network for densely connected local modules, which is analogous to searching for protein complexes in protein-protein interaction networks. We retained those significant local modules by comparing them with a negative control set. We then searched these significant modules for network cliques; those EKSes present in overlapping cliques are deemed as important for RBP binding. Analogous to the study of protein-protein interactions, this is akin to finding proteins that are “hub” nodes on a protein-protein interaction network.

We used the ClusterONE software (52) to derive densely connected modules; the parameters were set at (-s 100, -d 0.8, --max-overlap 0.2), i.e. requiring modules having density higher than 0.8 and a size of at least 100 nodes. The purpose of this step is to ensure the similarity network among EKSes we constructed is sufficiently dense to conduct maximal clique detections. To evaluate the significance of network modules of a specific RBP, we generated negative sites from non-target RBP binding datasets using a greedy strategy to ensure that both the positive set and the negative set have the similar single and di-nucleotide frequencies. (see **Supplementary Method S1** for details) We then compared Positional Weight Matrices (PWMs) of modules in the same structural context of these two sets by using Pearson *χ*^2^ test (53). We note that the *χ*^2^ p-value which is calculated for each column in PWM is based on the null hypothesis that the aligned columns are independent and identically distributed observations from the same multinomial distribution. We used the geometric mean of the column p-values as the p-value of module for the significance evaluation.

After confirming the existence of dense modules in the similarity network, we next applied Bron–Kerbosch algorithm (54-56) for maximal clique enumeration (MCE) over the similarity network to obtain the maximal cliques; we retained the upper half of the maximal cliques according to the rank of their sizes. A maximal clique is defined as a clique that cannot be extended by inclusion of additional adjacent vertices, i.e., it is not a subset of a larger clique. The objective of this step is to identify representative EKS or subgraph in a densely connected region in a similarity network so that it can capture the modality of the network. In practice, maximal cliques in the EKS similarity network are highly overlapping, in other words, an EKS can be part of multiple maximal cliques. These EKSes are likely representative of the overall topology of the dense subnetworks and the binding preference of the RBP, subsequently we extracted the top 50 of these recurring EKSes and considered them as RBP structural element that are important for RBP binding.

### 2.8 RNA 3D structure modeling and evaluation of network motifs

The RNA 3D structures were predicted using a discrete molecular dynamics (DMD) method (42) with predicted base-pair information provided as constraints. The DMD engine and scripts were adopted from the literature (57) and kindly provided by Dr. Feng Ding’s lab (http://dlab.clemson.edu/). Starting from an RNA sequence and base-pairing information, the default DMD settings were used to perform coarse-grained folding simulation for 100,000 DMD time units. For selected LIN28B systems, the last snapshots of DMD simulations were extracted to recover the 3-beads coarse-grained representation back to full atoms to perform more accurate full-atom docking and simulations. Since this conversion may result in severe clashes or abnormal bond length, some minimization and equilibration, as well as a short 10-ns full atom simulations were performed to further refine the structures before docking. For spatial accessibility, a 15Å cut-off was used to count the neighboring atoms of the given regions to calculate the exposure level, while nucleotides within 2 positions on both 5’ and 3’ directions were excluded since they would always be counted due to bonded connections except for at both ends of the sequences. To evaluate the significance of the discovered network motifs, Mann-Whitney-Wilcoxon (MWW) rank-sum U Test (58) was performed on such neighboring counts of identified motifs against the average of all the k-mer regions.

Charmm-gui webserver (59-61) was used to set up the MD systems, the complex was solvated in a water box with 15 A° buffer of water extending from the RNA or protein-RNA complex. K^+^ and Cl^2-^ ions were added to ensure a 0.15M ionic concentration and zero net charge. Due to the diversity and intrinsic flexibility of RNA molecules, the built systems contain a wide range of ∼300,000 to 700,000 atoms. After 10,000 steps of minimization and equilibration, where harmonic restraints were applied to heavy atoms, production MD simulations were performed in the NPT ensemble. The Nosé-Hoover method (62,63) was used with temperature T = 30°C. The Parrinello–Rahman method (64) was used for pressure coupling. A 10-Å switching distance and a 12-Å cutoff distance were used for non-bonded interactions. The particle mesh Ewald method (65) was used for electrostatics calculations. The LINCS algorithm (66) was used to constrain the hydrogen containing bond lengths for a 2-fs time step. The energy minimization and MD simulations were performed with the GROMACS program (67) version 2019.6-CPU/GPU using the CHARMM36m force field (68-70) and TIP3P water model (71).

The RNA structures after 10-ns production run were used as receptors and the LIN28B CSD structure (PDB ID 4A4I) (72) was used as ligand to perform protein-RNA docking using the ClusPro webserver (73-75); the predicted RNA motifs was provided as binding site constraints. The most populated docking poses were subjected to another 100 ns MD simulation to further test the binding stability; the last 50 ns equilibrated trajectories were used for analysis and visualization (**Figure S2**). We used the following geometric criteria to identify a hydrogen bond (HB) between two polar non-hydrogen atoms (i.e., acceptor and donor): the donor-acceptor distance is <3.5 Å, and the deviation of the donor-hydrogen-acceptor angle from 180° is <60°. We used the VMD program to identify and calculate the occupancy of HBs, the root-mean-square deviation (RMSD) of the protein and motif and for visualization (76).

## 3. RESULTS

### 3.1 Selection and optimization of *k* in GraphK partition

As shown in **Figure 1 (Step 2)** and **Figure 2**, the EKSes were constructed by traversing and extending *k* nucleotides from a specific base-pair. To select the most appropriate *k* to partition the RNA molecule, we investigated the occurrence of intervals between predicted base-pairs in the predicted RNA secondary structure.

We first divide all the nucleotides in the entire RNA as paired or unpaired and define the distance between two adjacent paired nucleotides as gap length. As shown in **Figure 3** (top panel), approximately 79.51% of all the gap lengths is 1, which represents stacked adjacent base pairs. The bottom of **Figure 3** shows, excluding gap length 1, gap length <= 9 can cover 87.91% of all possible gaps in 22 RBPs. Note that EKSes were constructed by extending from the initial base pair in two directions, thus a gap length of 9 is equivalent to 5-mer extension. This indicates that, with *k* ranging from 3 to 5, 87.91% of the nucleotides in RBP binding sites are likely covered by at least one EKS. We also note that most RNA binding domains tend to bind to short RNA sequences, therefore we investigated *k* up to 5 in our pipeline. The RNANetMotif pipeline is quite flexible so users can customize the cut-off *k* or other global parameters to suit their requirements. Moreover, the bottom bar plot in **Figure 3** shows that the number of occurrences decreases as the gap length increases.

**Figure 3.**
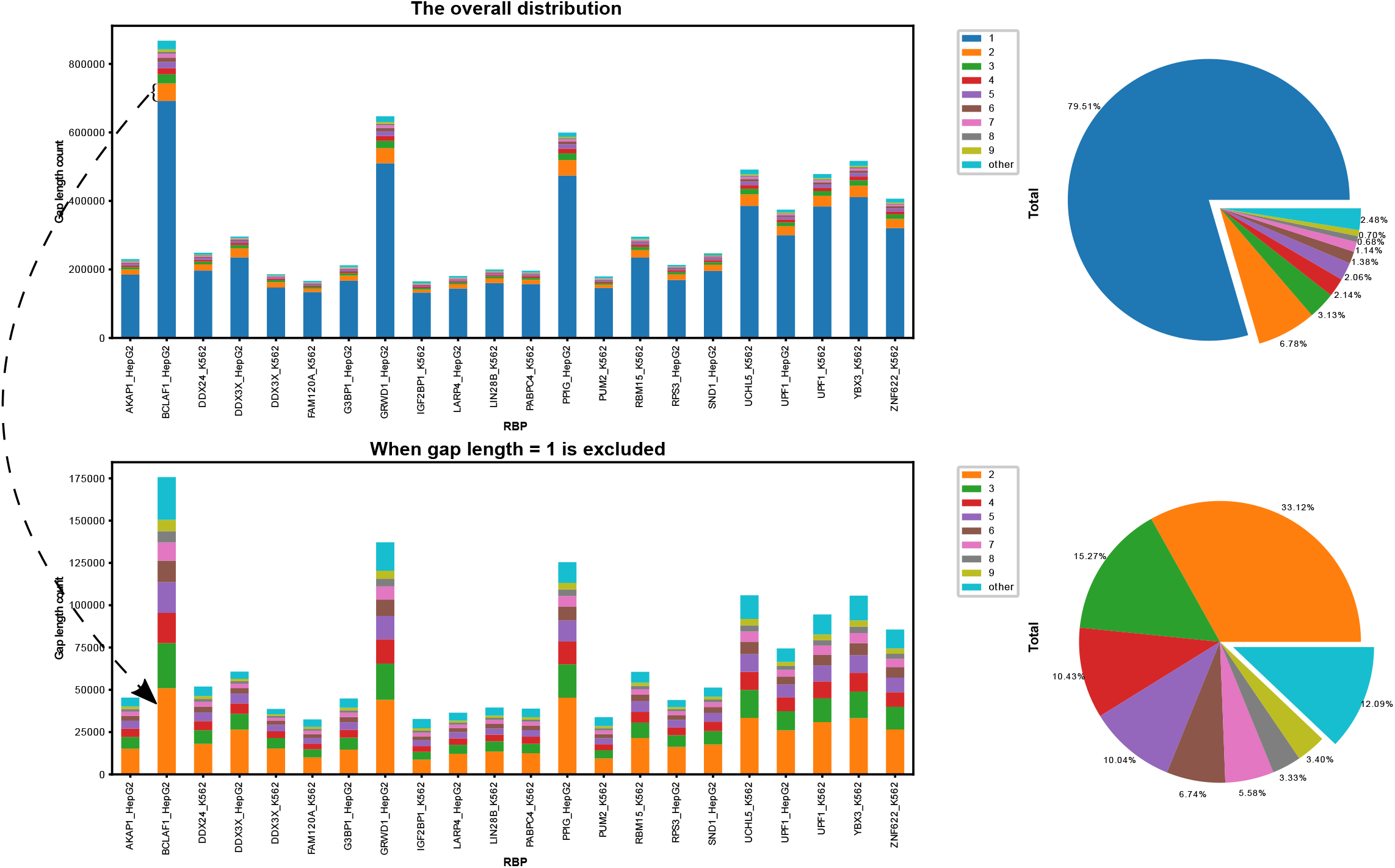
Distribution on the count of gap length of base-pairings.

After applying GraphK to partition the RBP binding sites into EKSes, we pooled these selected EKSes and investigated the properties of these elements. We further discretized the predicted secondary structure probability of each nucleotide into one of the following 5 states: P (paired), H (hairpin Loop), E (extended, unstructured), I (internal loop, bulge), M (multiple loop). We next counted the occurrence of sequence and structure of EKSes respectively; **Figure 4** shows the frequency of top sequence instances and top secondary structure instances in scatter plots for five selected RBPs on different *k*. From **Figure 4**, we find that after applying GraphK partition, the distribution of top sequence instances of EKS has lower frequencies than top structure instances. Given the low number of top sequence instances occurrence, we fitted a new distance and enrichment definition in our graph-preserving framework (**Method 2.6, 2.7**).

**Figure 4.**
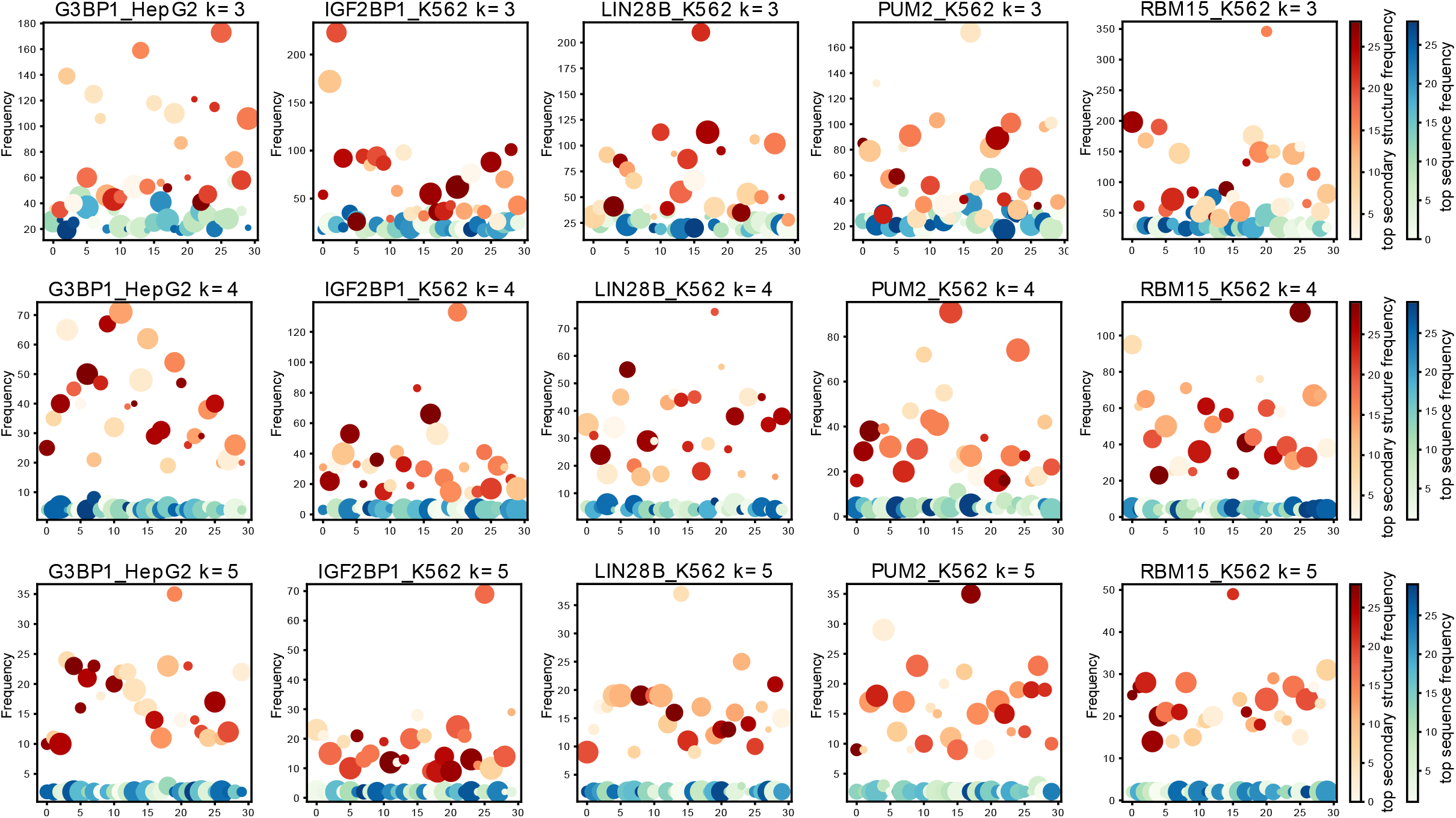
Distribution on the frequency of top sequence and structure instances in final EKS pool for 5 RBPs.

### 3.2 Significant modules in the similarity network of EKSes

We next calculated HVDM distances among pairs of EKSes and constructed a similarity network with individual EKS as nodes and HVDM distance as edge weight. We then used ClusterOne software to search for densely connected network modules in the similarity network and evaluate the statistical significance of these modules by Pearson *χ*^2^ test. As described above in section **3.1**, we next discretized secondary structure representation of each nucleotide and visualized the sequence-structure preference of these significant modules in **Figure 5** for 16 selected RBPs. We did not observe statistically significant modules for RBPs that preferentially bind to double-stranded RNAs such as DDX3X, DDX24, GRWD1 (containing WD repeat domain). We think this may be because RNA structure plays different roles in protein-RNA recognition for double-stranded RBPs than single-stranded RBPs, which is consistent with previous observations (40,77). We also did not observe significant EKS modules for two other RBPs PPIG, BCLAF1, which may be because these RBPs have multiple binding modes or multiple RNA binding domains, generating a mixture of signals (78). Further investigation is warranted for these RBPs.

**Figure 5.**
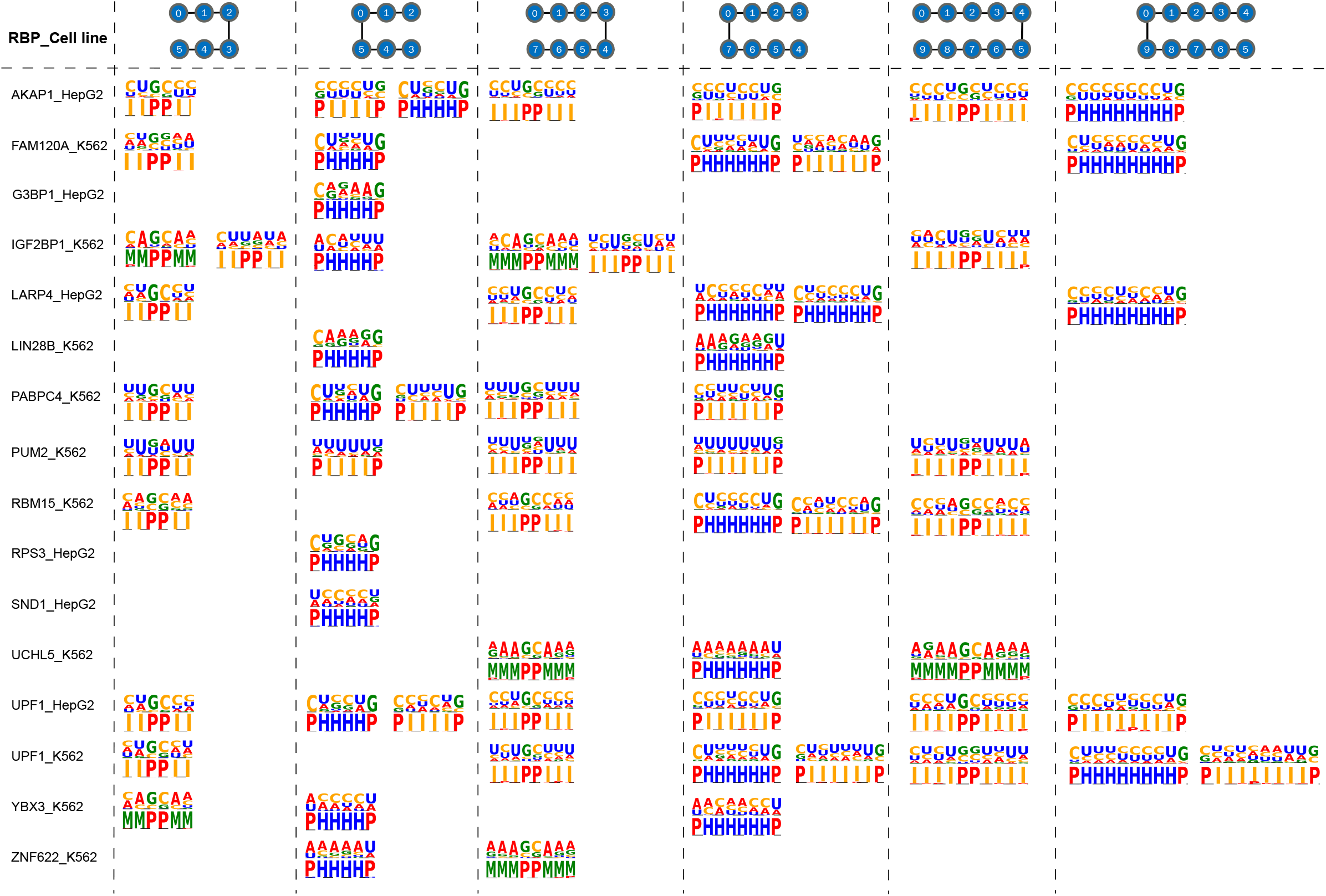
Combined sequence and structure logos of significant modules of 16 RBPs. Left-opened and right-opened EKSes of different sizes are displayed.

**Figure 5** shows the sequence logo and associated secondary structures of the enriched EKS modules for different k-mer sizes and EKS topologies. We notice that proteins with the same binding domains tend to have similar binding profiles. For example, IGF2BP1, PABPC4, LARP4 and RBM15 share a common RRM (RNA recognition Motif) domain, consisted of pyrimidine-rich internal loops and hairpin loops. Not surprisingly, we also note that the same proteins have similar profiles in different cell lines. For instance, profiles of UPF1 from HepG2 and K562 tend to share similar sequence preferences in both hairpin loops and internal loops when *k* is set at 5.

### 3.3 Significance of EKS motifs tested by 3D structure modelling

In the world of protein, linear polypeptides usually fold into certain three-dimensional structures to ultimately achieve cellular functions. Similarly, protein-RNA interactions often require RNA to fold into specific 3-D shapes in order to precisely spatially present relevant chemical moieties on sugar-phosphate backbone or nucleoside bases (79). It has been recognized in many cases that the complementarity in overall shape and structure between protein and RNA is as important as simple conservation of consensus RNA sequence. Just like in protein-protein interactions, in addition to static structures, intrinsic dynamics of RNA molecules also plays an important role in interaction between RNA and other molecules (80). Towards this goal, we performed large scale discrete molecular dynamics simulations to model the 3-D structure of four well-studied RNA binding proteins (G3BP1, RBM15, LIN28B, PUM2). In addition to the RNA dynamics study, for LIN28B, we also performed protein-RNA docking and all-atom molecular dynamics simulations to validate our predicted motifs.

Following the protocol described in **Methods (**section **2.8)**, we modeled the structure of bound RNA molecules for the four aforementioned RBPs. To achieve effective binding between protein and RNA, the relevant RNA moieties need to be exposed and accessible to amino acid residues on the RBP. We constructed coarse-grain structures of RNAs and calculated spatial accessibility of the predicted RNA structural motifs by counting the number of neighboring atoms within a 15 Å radius. Our hypothesis is that these structural motifs would have higher accessibility than the rest of the RNA molecules. For four well-studied RBPs (G3BP1, LIN28B, PUM2, RBM15), we performed one-sided Mann Whitney U Test (Wilcoxon Rank Sum Test) to compare the atom counts of the identified motif with all other k-mer regions. **Figure 6** shows that for all of these RBPs, the identified RNA structure motifs have statistically significant fewer atoms in their vicinity than the rest of the RNA molecule, indicating higher spatial accessibility for these motifs.

**Figure 6.**
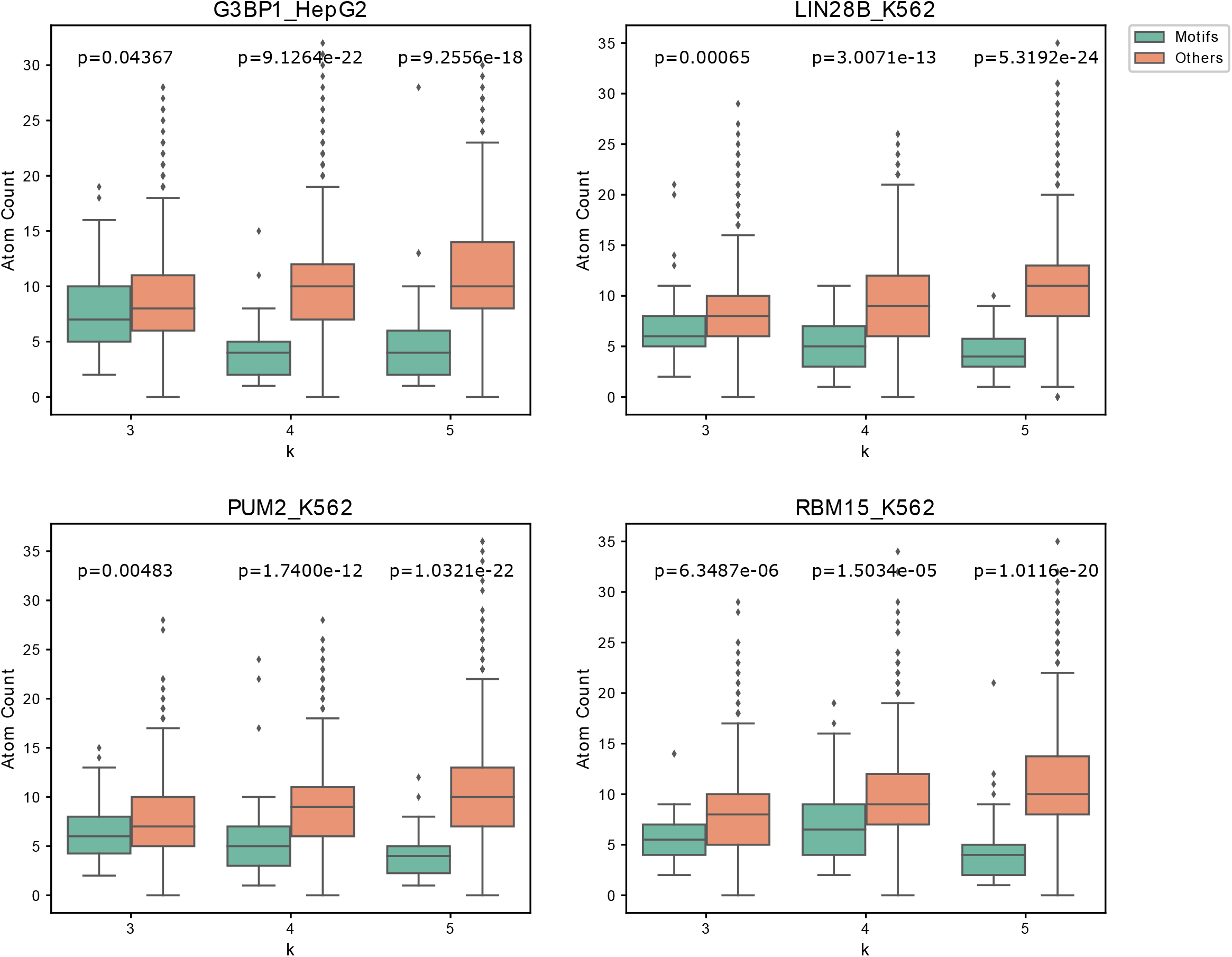
Boxplot of atom counts calculated from modeled 3D structure of motifs and other regions for 4 RBPs.

### 3.4 Case study: LIN28B-RNA docking and MD simulation

We next performed protein-RNA docking and simulation on LIN28 to further test the role of the predicted RNA structure motif in RBP-RNA recognition. We selected LIN28 for this purpose since this protein is well studied and there exist multiple high-resolution structure of it complexed with single stranded RNA (81,82).

We chose two representative sequences from the 5mer motifs and one from 4mer motifs to perform all-atom structure refinement and docking (see **Methods**). We chose the ClusPro web server (74) to perform RNA-protein docking, since it uses an FFT based method that can do a quick, accurate, and unbiased global search to explore protein-RNA binding modes. Interestingly, in all the tested cases, the binding poses with the lowest energy all captured the loop 4 in LIN28, which is a highly charged hairpin loop at the tip of the 4-5 beta sheet corresponding to residues 88-95. As shown in **Figure 7**, this loop is inserted into the motifs predicted by RNANetMotif. Subsequent MD simulations confirmed that this binding mode is energetically stable and strong phosphate contacts are formed while additional base contacts also formed, mainly through three Lys residues on this loop (Lys 88, 89, 92) (**Supplementary Table 2, Figure 7**). In the case of 4mer simulation, this loop-RNA contact served as an anchor and additional contacts were formed after certain RNA conformational changes. In a recent study, NMR and mutation studies indicated that K88, K89 (K98, K99 in LIN28A) play a role in electrostatic interactions with nucleic acids (83), and a recent study found a highly conserved Lys residue in YB1 protein (equivalent to K92 in Lin28B) is involved in ssDNA binding (84), as did the structural study of LIN28 on ssRNA nucleotides (72). Altogether, these evidences suggest that our graph-based approach is very effective in finding structure elements in RBP binding sites that are important in RBP-RNA recognition. To the best of our knowledge, such a unique approach has not been described in the literature.

**Table 2.**
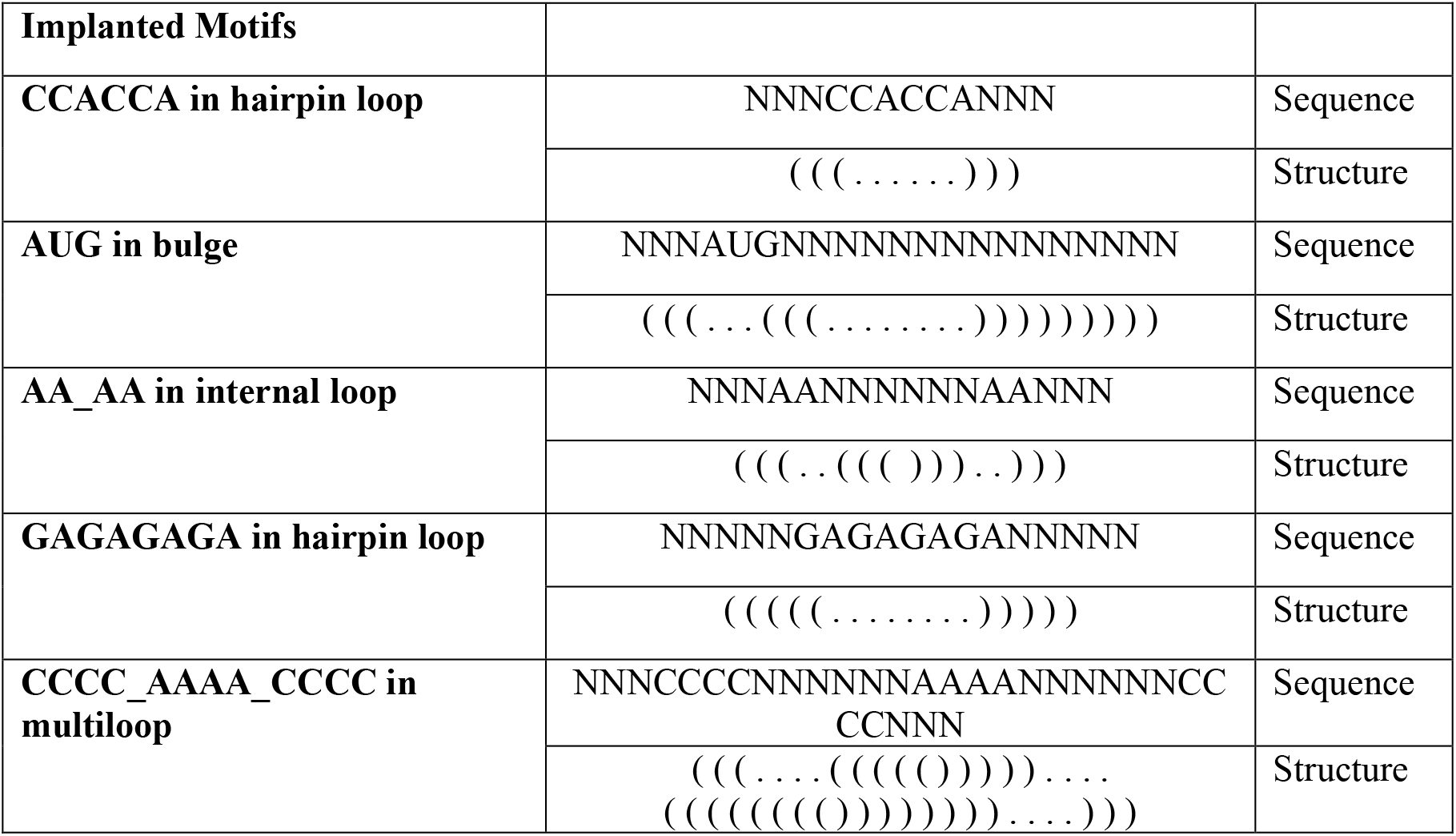
Implanted Motifs.

**Figure 7.**
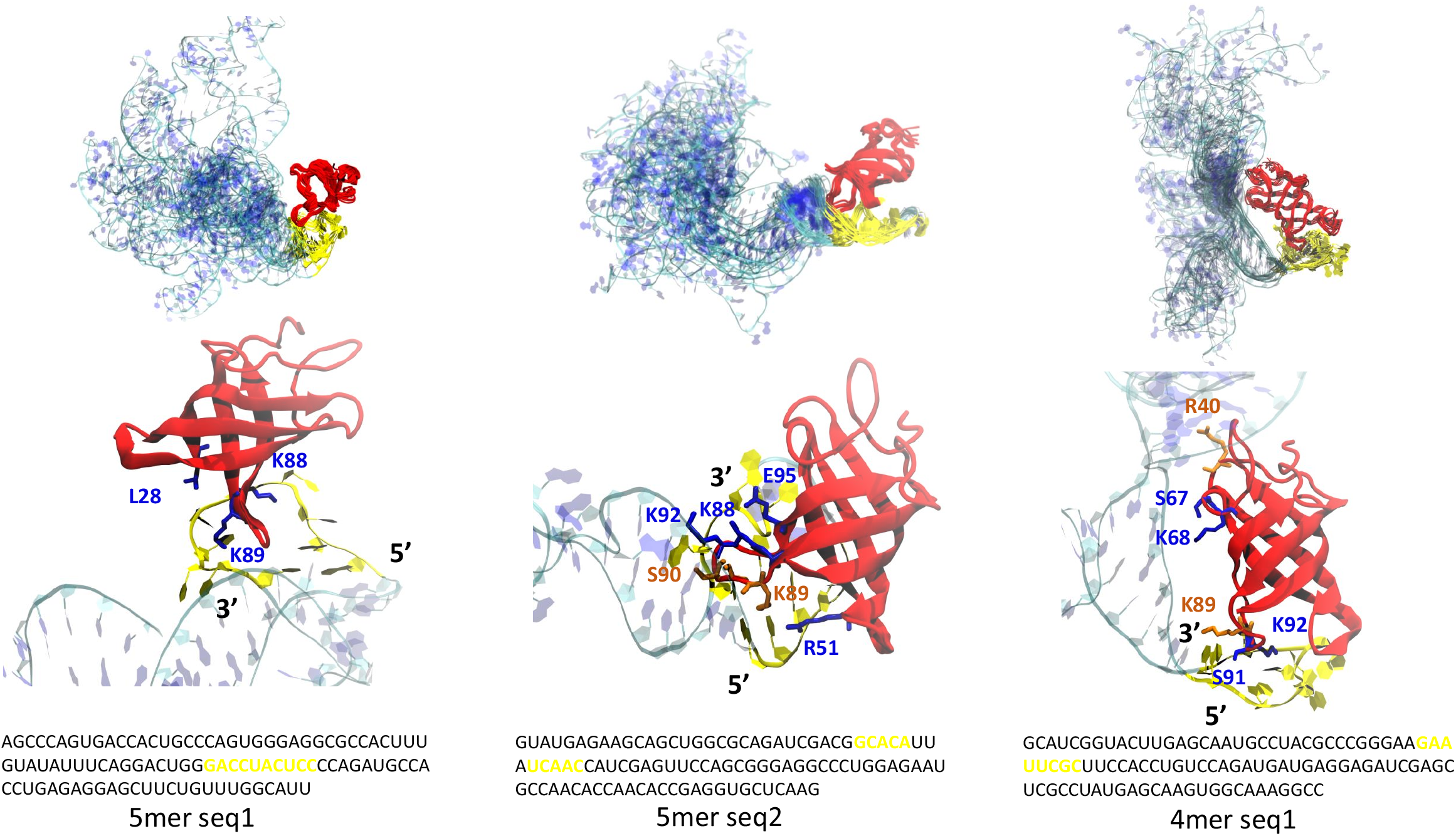
Complex simulation ensemble of docking of LIN28B CSD domain and three RNA network motifs. MD ensembles (top) and snapshots (bottom) showing the stability of Protein-RNA interface and key amino acids forming phosphate (blue) and base (orange) contact. LIN28B CSD protein is shown in red and 100 nt RNA shown in cyan and blue, identified motifs are shown in yellow with specific sequence information.

### 3.5 Comparison with other sequence-structure motif predictors on eCLIP datasets

We next compared RNANetMotif with three other computational methods, BEAM, GraphProt and ssHMM with two different secondary structure predictors on the eCLIP datasets. We displayed the motifs identified by RNANetMotif and by other methods in **Figure 8** and summarized the common motif elements in the right column. As seen in **Figure 8**, motifs identified by RNANetMotif closely resemble the motifs predicted by other methods on eCLIP datasets, for example, PUM2 recognizes U-rich internal loop and UCHL5 recognizes AGAA in the multiloop region.

**Figure 8.**
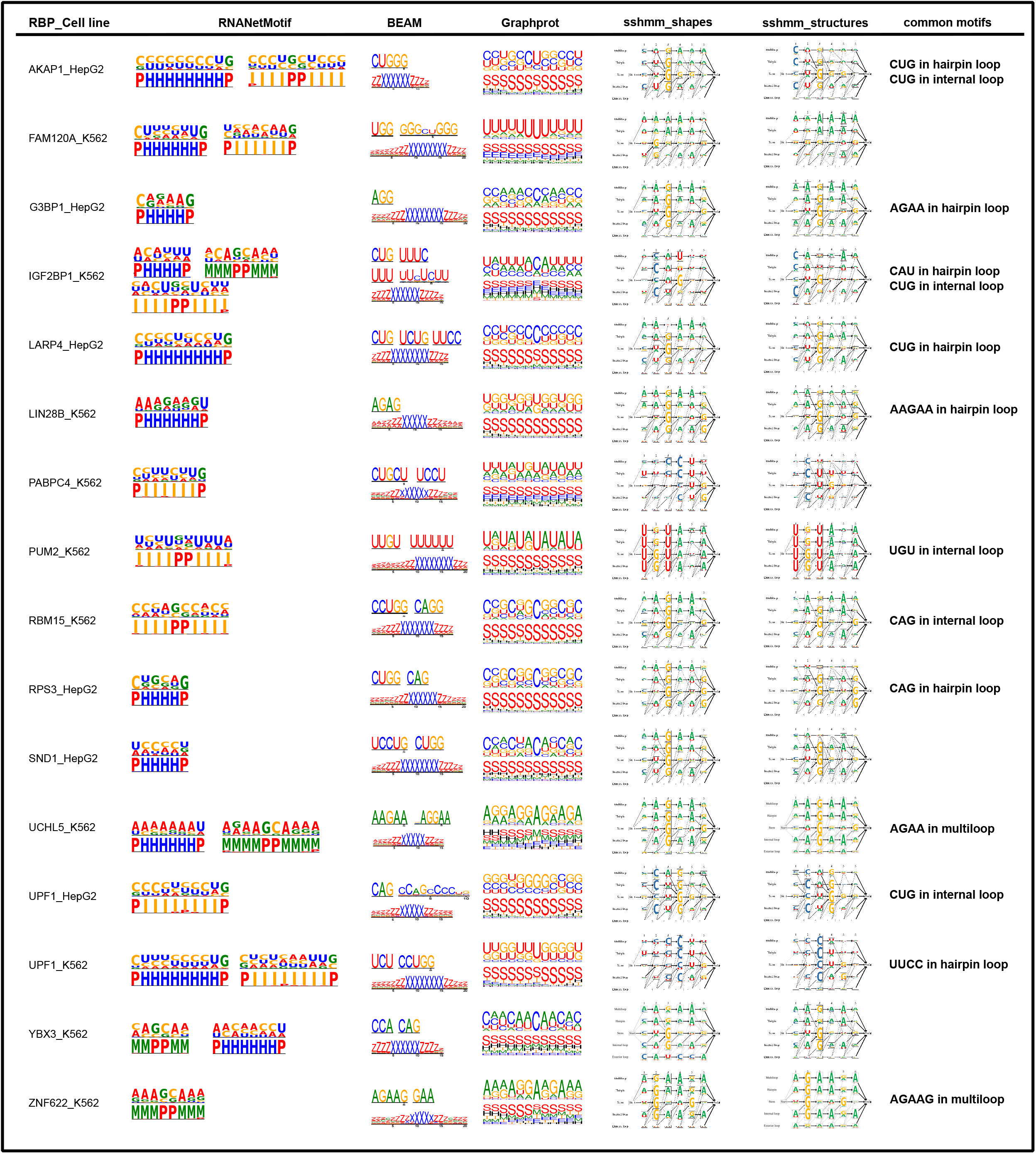
Comparison with other sequence-structure motif predictors on 16 eCLIP datasets.

### 3.6 Recovery of sequence and structure motifs from synthetic datasets

Having demonstrated the effectiveness of RNANetMotif in finding important RNA structural elements in RBP bound RNA sequences, we next investigated whether this approach is also effective in finding meaningful RNA structural motifs in a more general setting. As shown in **Table 2**, we selected five RNA sequence-structure motifs of representative types, i.e., hairpin loop, bulge, internal loop, hairpin loop and multi loop. We then randomized the flanking nucleotide sequences (represented as Ns) while maintaining the structure motif. We synthesized 100 instances of these sequence motifs and inserted each instance into a randomly generated RNA sequence of 100 nt long. For each type of motif, we also created a set of random background sequence set of the same size as the negative control set. We next ran RNANetMotif and three other methods, GraphProt, RNAcontext and Zagros, on these sequence sets to try to recover these spiked-in motifs. As shown in **Table 3**, RNANetMotif consistently recovered the sequence and structure of every one of the five motifs, which compares favorably with other methods. The detailed recovering results of these four methods are summarized in **Supplementary Materials**.

**Table 3.**
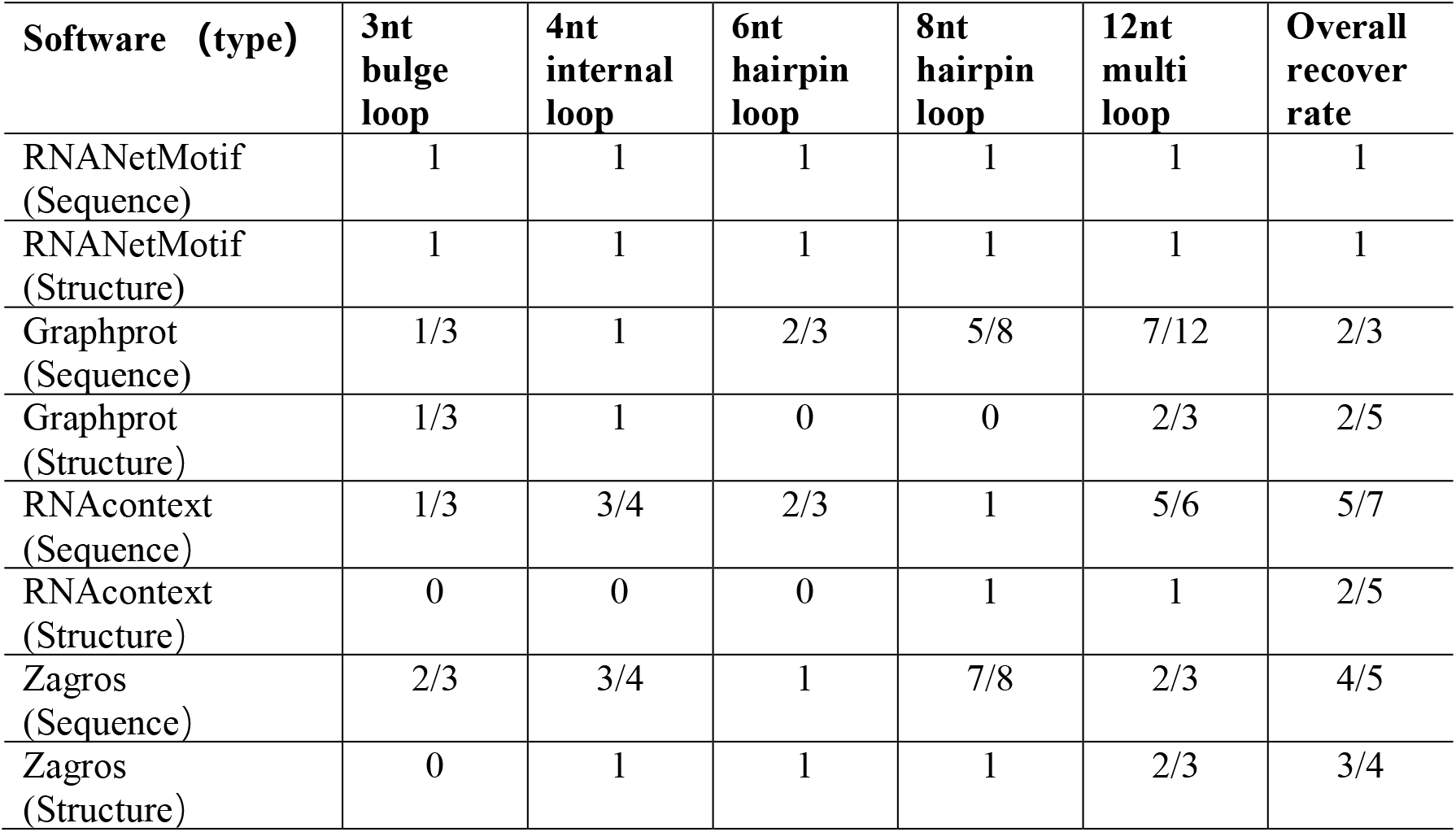
Recovery rate of sequence and structure on synthetic data for four methods

### 3.7 Run-time and memory comparisons

We compared the run time of RNANetMotif with other methods on five eCLIP datasets on a 32-core CPU (see Supplemental **Figure S3**). We excluded the time for RNA secondary structure prediction and model training time for Graphprot. RNANetMotif consumes the least time among all the five tools, while Graphprot ran a little slower than RNANetMotif on three datasets and faster than RNANetMotif on two other datasets. All the tools had similar level of memory usage, with maximum memory usage of no more than 10 GB on all the tested data. We like to note that a more comprehensive and robust investigation and benchmark is warranted to test the utility of RNANetMotif with regard to this task.

## 4. DISCUSSION

In this paper we describe a novel network-based approach in finding meaningful sequence and structure motifs from RBP bound RNA sequences. We recognize that there have been many other methods developed over the years that have addressed this problem from many different angles (reviewed in **Introduction)**. The novelty of RNANetMotif is that it takes a graph-based approach to extract enriched subgraphs from RNA secondary structures. Ideally, the most accurate and unbiased way to determine the binding mechanism between an RBP and its target RNAs is to compare high-resolution 3-dimensional structures of a representative set of RBP-RNA complexes and derive a set of common 3-D structure elements shared by these structures. However, this is technically challenging and logistically unrealistic. It has been well documented that linear representations such as Positional Weight Matrices (PWM) have their limitations in capturing the binding preferences (85,86). A number of methods such as BEAM or GraphProt have improved upon PWMs by adding structural descriptors to each position, i.e., helix, stem, loop, etc, which has showed improvement (16,25). Despite these recent developments, we feel there is still room for improvement. Motivated by the observation that RNA secondary structures, represented as a network, are essentially low-resolution abstracted structure of RNA molecules, we hypothesized that there exist enriched subgraphs in these networks that are determinants of the recognition process between RBP and their RNA targets. Conceptually, this is analogous to the study and design of protein structures from the aspect of hydrogen bond and other interactions among amino acid residues.

We introduced a new concept to represent RNA elements, i.e., EKS (Extended K-mer Subgraphs) and an algorithm GraphK. As demonstrated in this study, we think EKS is a promising approach in mining RNA secondary structures for enriched motifs. We further showed in **Result 3.6** that it can be extended to the study of more general problems in addition to RBP-RNA interactions, although more rigorous benchmarking is required. Compared with other methods, RNANetMotif has the following unique characteristics. (i) It recognizes RNA target sequences as a network and applies graph theory approaches to extract meaningful and enriched subgraphs. (ii) In contrast to other methods, RNANetMotif follows an unsupervised scheme thus avoids the noise and uncertainty introduced by the false negative data. (iii) We also conducted structure modeling and molecular dynamics studies and validated some of the predictions. The RNANetMotif is a new way of investigating RNA sequences and can be further extended into the analysis of other categories of RNA sequences. The source code of RNANetMotif can be accessed at Github (https://github.com/hongli-ma/RNANetMotif). We also constructed a website that allow users to test their own data: http://rnanetmotif.ccbr.utoronto.ca.

There are several limitations and avenues for future approvement in our current approach. When building a network of RNA secondary structure interactions, we only included canonical Watson–Crick and GU wobble base pairs and excluded other non-canonical base pairing such as Hoogsteen base pairing or Leontis-Westhof interactions (87,88). This is mostly due to the lack of high-resolution 3-D RNA structures, which makes accurate prediction of these non-canonical interactions less feasible. We anticipate deep learning models, especially graph models, can probably alleviate this challenge in the coming years(33,89-91). We also focused on in vivo RBP binding data in this study; it would be very interesting to extend this graph-based approach to in vitro data generated by RNAcompete or HTR-SELEX.

## 5. DATA AVAILABILITY

The RNANetMotif software is available for download at https://github.com/hongli-ma/RNANetMotif. eCLIP datasets used in this study can be found at “Download” page on our webserver (http://rnanetmotif.ccbr.utoronto.ca).

## 7. ACKNOWLEDGEMENTS

This research was enabled in part by support provided by Compute Canada (www.computecanada.ca). Computations were performed on the Niagara supercomputer at the SciNet HPC Consortium. SciNet is funded by: the Canada Foundation for Innovation; the Government of Ontario; Ontario Research Fund - Research Excellence; and the University of Toronto. HM thanks Shandong University for their financial support through scholarships under overseas exchange program. HW would like to thank Dr. Philip Kim from CCBR, University of Toronto for post-doctoral fellowship support in the early stage of this work. We thank Dr. Feng Ding and Zhenzhen Zhang from Clemson University for their support and useful discussion about DMD method.

## 8. AUTHOR CONTRIBUTIONS

HM and ZZ conceived the project. HM collected the data ad developed the algorithm.

HW conducted molecular dynamics and modeling. ZX implemented the webserver.

HM, HW and ZZ wrote the paper. GL and ZZ supervised the project.

