## Supplemental Methods and figures for "RNANetMotif: identifying sequence-structure RNA network motifs in RNA-protein binding sites"

### Supplementary Methods

#### S1: Selection of control sites for the evaluations of network modules

To evaluate the significance of network modules of a specific RBP, we selected negative sites from non-target RBP binding sites' sets as a control set. To ensure that the negative set of each RBP has similar nucleotide frequencies as the corresponding positive set, we adopted a greedy strategy. The detailed steps are as follows.

Denote the target RBP as P, other 21 RBPs as OP. And the set containing the binding sites of P is denoted as $S_P$, binding sites of OP as $S_{OP}$. Each 100nt site s in $S_P$ and $S_{OP}$ is mapped to a 4D vector $v_s = (v_{sA}, v_{sC}, v_{sG}, v_{sU})$, where $v_{st}, t \in \{A, C, G, U\}$ is the frequency of t in s. And A%, C%, G%, U% distribution in binding sites of every RBP is displayed in **Figure S1**.

Define the Chebyshev distance between two sites $s_i$ and $s_j$ as

$$D_{Chebyshev}(s_i, s_j) = D_{Chebyshev}\left(v_{s_i}, v_{s_j}\right) = max_{n\in\{A,C,G,U\}}(|v_{s_i n} - v_{s_j n}|)$$

**Algorithm 1.** Greedy algorithm

**procedure** GREEDY $(S_P, S_{OP})$
    index ← 0
    N ← { }
    **while** |N|<$|S_P|$ **do**
        next_neg ← $min_{S_{OP}} D_{Chebyshev}(S_P(index),\ c), \forall\ c \in\ S_{OP} \backslash N$
        N ← N ∪ next_neg
        index ← index + 1
    **end while**
    pValue ← KS_test($S_P$, N)
    **if** pValue ≤ ε **then**
        **return** N
    **end if**
**end procedure**

We then removed redundant and missing data for control sets of each RBP using module 'cd-hit-est' from CD-HIT at 80% similarity. For each RBP, negative sites with sequence similarity above 80% to any positive sites were excluded using module 'cd-hit-est-2d' from the CD-HIT tool.

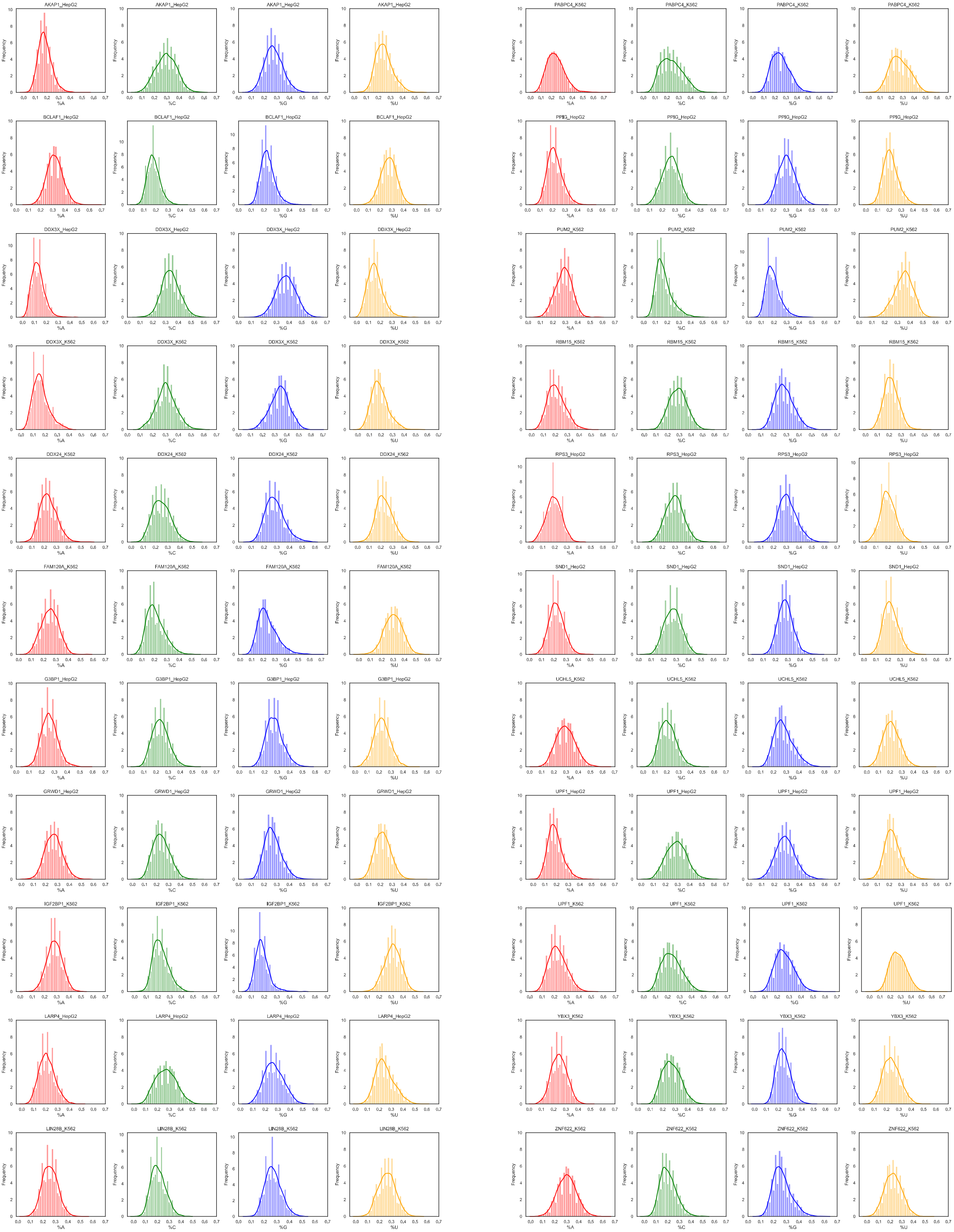


**Figure S1.** Nucleotide distribution in binding sites of 22 RBPs.

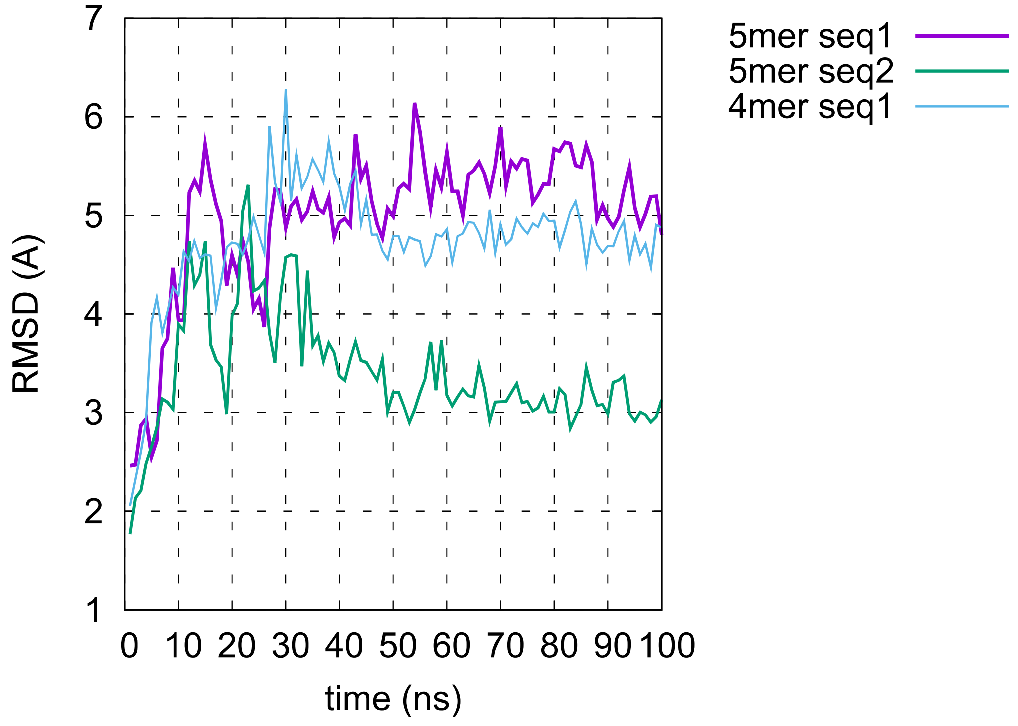


**Figure S2.** RMSD plot of molecular dynamics trajectories.

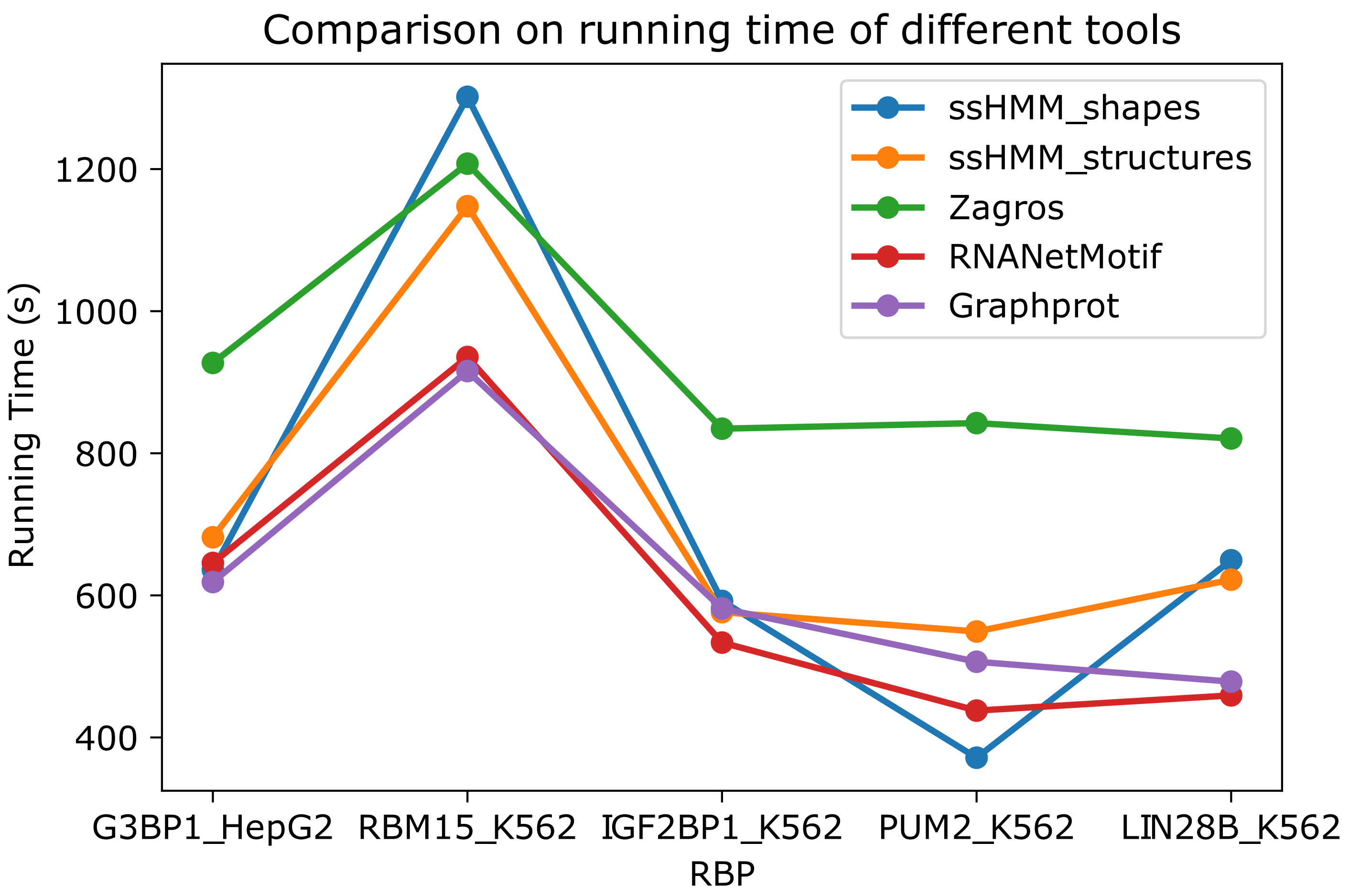


**Figure S3.** Comparison on running time of different tools on 5 eCLIP datasets.
